# The Interaction of Genotype and Environment Determines Variation in the Maize Kernel Ionome

**DOI:** 10.1101/048173

**Authors:** Alexandra Asaro, Greg Ziegler, Cathrine Ziyomo, Owen A. Hoekenga, Brian P. Dilkes, Ivan Baxter

## Abstract

Plants obtain soil-resident elements that support growth and metabolism via water-mediated flow facilitated by transpiration and active transport processes. The availability of elements in the environment interact with the genetic capacity of organisms to modulate element uptake through plastic adaptive responses, such as homeostasis. These interactions should cause the elemental contents of plants to vary such that the effects of genetic polymorphisms influencing elemental accumulation will be dramatically dependent on the environment in which the plant is grown. To investigate genotype by environment interactions underlying elemental accumulation, we analyzed levels of elements in maize kernels of the Intermated B73 x Mo17 (IBM) recombinant inbred population grown in 10 different environments spanning a total of six locations and five different years. In analyses conducted separately for each environment, we identified a total of 79 quantitative trait loci controlling seed elemental accumulation. While a set of these QTL were found in multiple environments, the majority were specific to a single environment, suggesting the presence of genetic by environment interactions. To specifically identify and quantify QTL by environment interactions (QEIs), we implemented two methods: linear modeling with environmental covariates and QTL analysis on trait differences between growouts. With these approaches, we found several instances of QEI, indicating that elemental profiles are highly heritable, interrelated, and responsive to the environment.

**Author Summary:** Plants take up elements from the soil, a process that is highly regulated by the plant’s genome. To investigate how maize alters its elemental uptake in response to different environments, we analyzed the kernel elemental content of a population derived from a cross grown 10 different times in six locations. We found that environment had a profound effect on which genetic loci were important for elemental accumulation in the kernel. Our results suggest that to have a full understanding of elemental accumulation in maize kernels and other food crops, we will need to understand the interactions identified here at the level of the genes and the environmental variables that contribute to loading essential nutrients into seeds.

## Introduction

The intake, transport, and storage of elements are key processes underlying plant growth and survival. A plant must balance mineral levels to prevent accumulation of toxic concentrations of elements while taking up essential elements for growth. Food crops must strike similar balances to provide healthy nutrient contents of edible tissues. Adaptation to variation in soil, water, and temperature requires that plant genomes encode flexible regulation of mineral physiology to achieve homeostasis (1). This regulation must be responsive both to the availability of each regulated element in the environment and to the levels of these elements at the sites of use within the plant. Understanding how the genome encodes responses to element limitation or toxic excess in nutrient-poor or contaminated soils will help sustain our rapidly growing human population (2).

The concentrations of elements in a plant sample provide a useful read-out for the environmental, genetic and physiological processes important for plant adaptation. We and others developed high-throughput and inexpensive pipelines to detect and quantitate 20 different elemental concentrations by inductively coupled plasma mass spectrometry (ICP-MS). This process, termed *ionomics*, is the quantitative study of the complete set of mineral nutrients and trace elements in an organism (its *ionome*) (3). In crop plants such as maize and soybean, seed element profiles make an ideal study tissue as seeds provide a read-out of physiological status of the plant and are the food source.

Quantitative genetics using structured recombinant inbred populations is a powerful tool for dissecting the factors underlying elemental accumulation and relationships. By breaking up linkage blocks through recombination and then fixing these new haplotypes of diverse loci into mosaic sets of lines, these populations allow similar sets of alleles to be repeatedly tested in diverse environments (4). A variety of quantitative statistical approaches can then be used to identify QTL by environment interactions (QEI).

Here, we used elemental profiling of a maize recombinant inbred population grown in multiple environments to identify QTL and QEI underlying elemental accumulation. We sought both environmental and genetic determinants by implementing single-environment QTL mapping and analyses of combined data from multiple environments. Overall, we detected 79 loci controlling elemental accumulation, many of which were environment-specific, and identified loci exhibiting significant QEI.

## Results

### Genetic Regulation of Elemental Traits

The data used for this study is comprised of 20 elements measured in the seeds from *Zea mays* L. Intermated B73 x Mo17 recombinant inbred line (IBM) populations grown in 10 different location/year settings. The IBM population is a widely studied maize population of 302 intermated recombinant inbred lines, each of which have been genotyped with a set of 4,217 biallelic single nucleotide polymorphism (SNP) genetic markers (5). The four rounds of intermating and subsequent inbreeding resulted in more recombination and a longer genetic map for the IBM than for typical biparental recombinant inbred line populations. The number of individuals, marker density, and greater recombination facilitates more precise QTL localization than a standard RIL population (6–11). This greater resolution reduces the number of genes within a QTL support interval and increases the utility of QTL mapping as a hypothesis test for shared genetic regulation of multiple traits and aids in the discovery of the molecular identity of genes affecting QTL. For this study, subsets of the IBM population were grown at Homestead, Florida in 2005 (FL05) and 2006 (FL06), West Lafayette, Indiana in 2009 (IN09) and 2010 (IN10), Clayton, North Carolina in 2006 (NC06), Poplar Ridge, New York in 2005 (NY05), 2006 (NY06), and 2012 (NY12), Columbia, Missouri in 2006 (MO06), and Limpopo, South Africa in 2010 (SA10) (Table S1). While very few of the 233 unique IBM lines in the experiment were grown in all environments, 106 of the 233 lines were grown in 7 or more environments and 199 were grown in 3 or more environments. Within each growout, all samples were treated identically: seeds from all environments were stored in temperature and humidity controlled storage rooms after harvest and then shipped to the ionomics lab. We do not expect any change in ion composition from storage within a growout, however we cannot rule out that some of the differences between growouts might be due to slightly different moisture content. These differences are not likely to account for the genetic by environment interactions we observe as they should have similar effects on all lines. Single seeds were profiled for the quantities of 20 elements using ICP-MS and these measurements were normalized to seed weight and technical sources of variation using a linear model with the resulting values used as the elemental traits for all analysis (12). After outlier removal, seed element phenotypes were derived by averaging line replicates (kernels subsampled out of pooled ears from a row) within an environment.

Variation in the elemental traits was affected by both environment and genotype. Elemental traits generally exhibited lower heritability among genotypes grown across multiple environments than among genotype replicates within a single environment (Table 1). The broad-sense heritability (H2) of seed weight, 15 of 21 elements in NY05, 13 of 21 elements in NC06,and 13 of 21 elements in MO06 exceeded 0.60. Elements exhibiting low heritability within environments corresponded to the elements that are prone to analytical artifacts or present near the limits of detection by our methods, such as B, Al, and As. Seven elements had a broad sense heritability of at least 0.45 in a single environment (NY05, NC06, and NY06) but less than 0.1 across all environments. This decrease in heritability across the experiment, which was particularly striking for Mg, P, S, and Ni, is consistent with strong genotype by environment interactions governing the accumulation of these elements.

**Table 1.** Broad-sense heritability (H2) of element concentrations.

| **Trait** | **All**<br>**env** | **NY05** | **NC06** | **MO06** |
| --- | --- | --- | --- | --- |
| Seed<br>Weight | 0.51 | 0.59 | 0.69 | 0.89 |
| B | 0.02 | 0.35 | 0.51 | 0.06 |
| Na | 0.11 | 0.34 | 0.23 | 0.19 |
| Mg | 0.04 | 0.77 | 0.69 | 0.75 |
| Al | 0.10 | 0.39 | 0.50 | 0.08 |
| P | 0.04 | 0.62 | 0.69 | 0.33 |
| S | 0.05 | 0.73 | 0.77 | 0.51 |
| K | 0.07 | 0.69 | 0.72 | 0.36 |
| Ca | 0.15 | 0.65 | 0.63 | 0.77 |
| Mn | 0.16 | 0.80 | 0.80 | 0.75 |
| Fe | 0.07 | 0.76 | 0.73 | 0.63 |
| Co | 0.08 | 0.65 | 0.54 | 0.42 |
| Ni | 0.06 | 0.84 | 0.54 | 0.82 |
| Cu | 0.20 | 0.80 | 0.75 | 0.92 |
| Zn | 0.07 | 0.68 | 0.73 | 0.86 |
| As | 0.02 | 0.37 | 0.45 | 0.01 |
| Se | 0.04 | 0.32 | 0.35 | 0.68 |
| Rb | 0.03 | 0.49 | 0.45 | 0.69 |
| Sr | 0.07 | 0.61 | 0.48 | 0.53 |
| Mo | 0.29 | 0.85 | 0.73 | 0.96 |
| Cd | 0.55 | 0.71 | 0.69 | 0.24 |

All env: Line replicate averages from each location
NY05: 50 lines with 2 reps, 199 lines with 3 reps
NC06: 121 lines with 2 reps, 53 lines with 3 reps, 4 lines with 4 reps
MO06: 50 lines with 2 reps, 18 lines with 3 reps
\*outliers for each element calculated with outlier removal function, designated as NA \*for each single environment, for each trait, only lines w/o missing data and with reps >1 used to calculate heritability

A stepwise algorithm, implemented via *stepwiseqtl* in the R package R/qtl (13), was used to map QTL for seed weight and 20 seed elemental phenotypes. The stepwise algorithm iterates through the genome and tests for significant allelic effects for each marker on a phenotype. Forward and backward regression was used to generate final genome-wide QTL models for each trait. This QTL mapping procedure was completed for each of the IBM populations from each of the 10 environments for all 21 traits as separate analyses. QTL significance were determined using the 95th percentile threshold from 1000 *scanone* permutations as a penalty score for adding QTL to the stepwise model (14).

The environmental dependence on QTL detection was first estimated by identifying QTL common to multiple environments. If QTL detected in two or more growouts affected the same element and localized less than 5 cM apart they were considered to be the same locus. Across the 10 environments, a total of 79 QTL were identified for seed weight and 18 of the 20 elemental traits tested (none for Al or Co) (Fig 1B &C). Of these QTL, 63 were detected in a single environment and 16 were detected in multiple environments. The 16 QTL found in multiple environments included QTL detected in nearly all of the environments and QTL detected in only two. One QTL for Mo accumulation, on chromosome 1 in the genetic region containing the maize ortholog of the Arabidopsis molybdenum transporter MOT1 (15), was found in nine environments (Fig 1A). Another QTL affecting Cd accumulation, on chromosome 2 and without a clear candidate gene, was found in eight environments. Other QTL were only present in a smaller set of environments, such as the QTL for Ni accumulation on chromosome 9, which was found in five environments (Fig 1D). The strength of association and percent variance explained showed strong differences between environments even for these QTL that were detected in multiple environments (Table S2).

**Figure 1.**
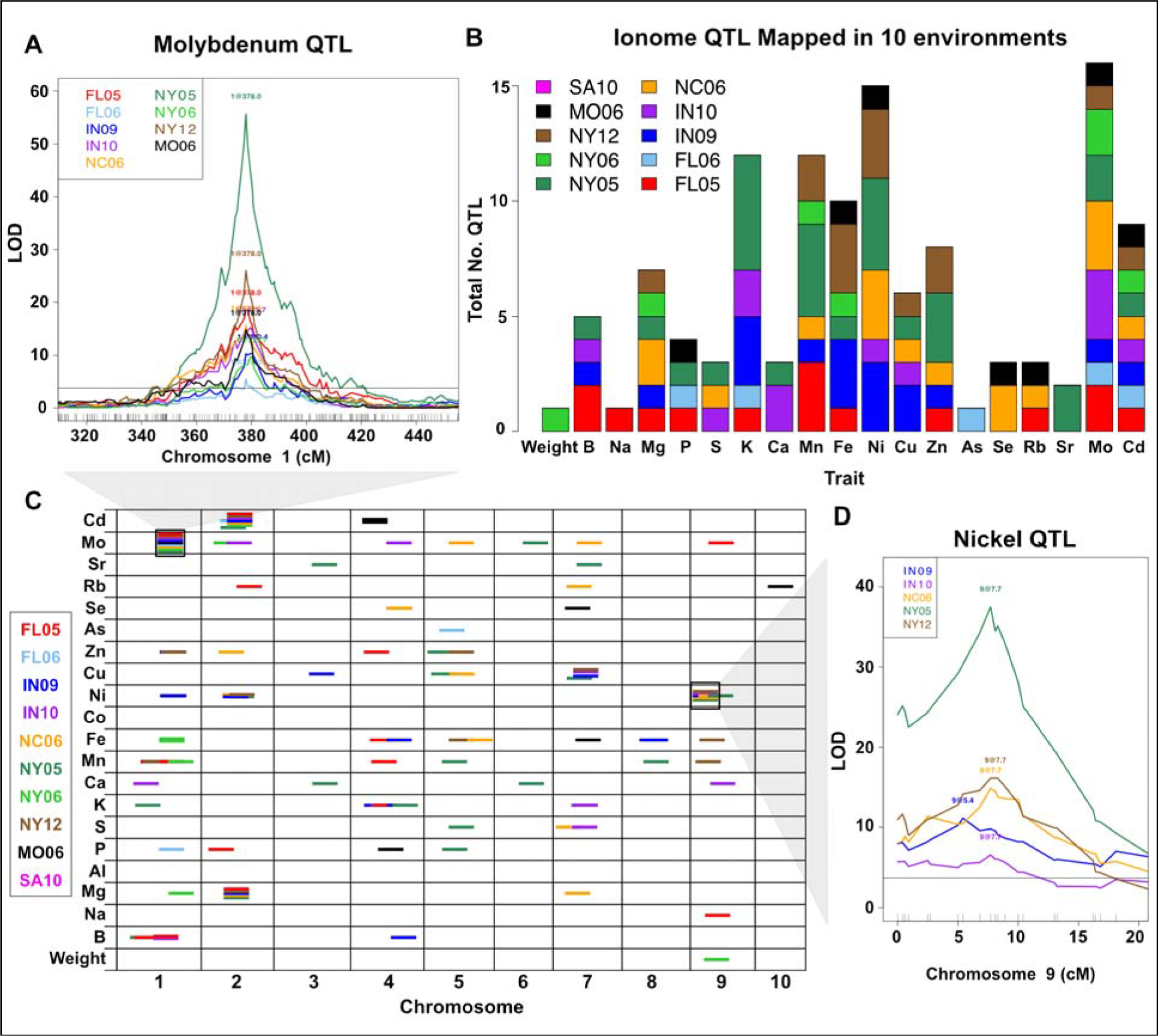
Ionome QTL from 10 Environments. QTL identified for seed weight and 20 element accumulation traits using the B73 x M017 intermated RIL population grown in 10 environments. (A) QTL on chromosome 1 affecting variation in molybdenum accumulation. An interval of Chr1 is shown on the x-axis (in centi-Morgans). The LOD score for the trait-genotype association is shown on the y-axis. The horizontal line is a significance threshold from 1000 random permutations (α = 0.05). The LOD profiles are plotted for all environments in which the highlighted QTL was detected. (B) Total number of QTL detected for each trait, colored by environment. (C) Significant QTL (α = 0.05) for each trait. QTL location is shown across the 10 maize chromosomes (in cM) on the x-axis. Dashes indicate QTL, with environment in which QTL was found designated by color. All dashes are the same length for visibility. The two black boxes around dashes correspond to LOD profiles traces in (A) and (D). (D) Stepwise QTL mapping output for nickel on chromosome 9. LOD profiles are plotted for all environments in which the QTL is significant.

As seen in the full-genome view of all QTL colored by environment (Fig 1C), there is a high incidence of QTL found in single locations. There are three hypotheses that could explain the large proportion of QTL found only in a single location: 1) strong QTL by environment interaction effects, 2) false positive detection of a QTL in an individual location and 3) false negatives assessment of QTL absence due to genetic action but statistical assessment below the permutation threshold in other locations. To reduce the risk of false positives in a single environment’s QTL set, the significance threshold was raised to the 99th percentile, where 31 of the 63 location-specific QTL remained significant. Despite the large number of trait/environment combinations tested (20 traits in 10 environments), the number of QTLs detected is much larger than the null expectation derived from a Bonferroni correction: 10 QTL (95th percentile threshold) and two QTL (99th percentile threshold). To account for false negatives, we scanned for QTL using a more permissive 75th percentile cutoff. Of the 63 single-environment QTL, only nine had QTL in other environments by this more permissive threshold. Thus, the majority of the 63 single-environment QTL most likely result from environmentally contingent genetic effects on the ionome.

#### QTL by Environment Interactions

That QTL detection was so strongly affected by environment suggested that the effects of allelic variation on element concentration were heavily dependent on environmental variables. These results, however, did not specifically test for QTL by environment interactions (QEI). Comparison between environments in our data is additionally complicated because different subsamples of the IBM population were grown at these different locations and years. While there are many different approaches to identifying QEI described in the literature (summarized in El-Soda et al. (16)) we focused on two previously implemented methods. The first considered location (but not year) by comparing the goodness of fit for linear models with and without an interactive covariate (17–19). The second method takes advantage of the ability to grow the same RILs in multiple years. Trait values measured in the same IBM line for the same element at the same site but in different years were subtracted from each other and the difference between years was assigned as the trait value for that RIL genotype for QTL detection (20, 21).

### Linear model estimation of QTL by location effects

The most common approach to analyze QEI is to fit a linear model with environment as both a cofactor and an interactive covariate and compare results to a model with environment as an additive covariate (22). This method is most amenable when data are available for the same lines grown in every environment, which was not the case across all of our dataset. Data from the three locations with two replicate years each (FL, IN, NY) were analyzed to reduce the number of covariates and increase the power to detect variation from the environment. The data for both years in each location were combined (FL05 & FL06, IN09 & IN10, NY05, NY06 & NY12), designating covariates based on location.

Two linear QTL models were fit to the combined data using the FL, IN, and NY locations as covariates. These models, as detailed in Bhatia et al., reflect the dependence of phenotypic variance on genetic variance, environmental variance, and genetic by environmental variance.

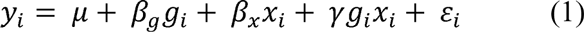

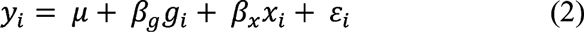

The first equation fit (1) is the full model considering the phenotype of individual *i* (*y_i_*) as controlled by genotype (*g_i_*), location (*x_i_*), and genotype by location interaction (*g_i_x_i_*), while the reduced model (2) estimates phenotypic without considering genotype by location interaction, using genotype and location as purely additive factors. *B_g_* and *B_x_* represent the additive effects of genotype and environment, respectively, while γ represents the effect of the genotype by environment interaction. These effects can be estimated through comparison of likelihood functions for each model to a null model. Subtracting the likelihood ratios of the reduced model (2) from the full model (1) will isolate the effect of genetic by environment interaction.

The program R/qtl was used to fit QTL using both the full and reduced models for sample weight and 20 elements, with three locations encoded as covariates in the environment term. For each marker, LOD scores resulting from the reduced QTL model were subtracted from LOD scores determined by the full model, leaving a LOD score for each marker representing solely the significance of the genetic by location component. The significance threshold for the subtracted LOD scores was calculated by using 1000 permutations of the three step procedure (fitting the two models with randomized data and then subtracting LOD scores). Even with this underpowered dataset, 10 QTL by location interactions exceeded the threshold (α = 0.05, Table 2). Interactions between QTL and location are likely to be due to a combination of soil and weather differences across different locations. In the case of Ni, our initial single-element QTL mapping conducted separately on data from each environment identified differences in QTL presence or strength between FL, IN, and NY locations for a QTL located at the beginning of chromosome 9 (Fig 2). This QTL corresponds to a locus found to have a significant QTL by location effect (Table 2). Remarkably, all elemental QTL by location interactions detected by this approach affected trace element accumulation. These elements are both low in concentration in the grain, and often variable among soils (23). Cd, an element for which we found significant QEI, has detrimental effects on both human and plant health (24) and is toxic in food at levels as low as .05 ppm. (25). The locus with the strongest QEI for Cd does not follow location averages of Cd content in the grain (Table S3) and therefore is unlikely to be affected by crossing a detection threshold driven by higher Cd in the soils at those locations. The lack of direct correlation between QTL significance and grain content also occurs for the loci with strong by-location effects for Mo and Ni. This demonstrates that reduced cadmium or enhanced micronutrient contents in grain require plant breeding selections that consider complex genetic by environment interactions rather than genotypes assessed in a single soil environment.

**Table 2.** QTL with Significant by-Location interactions.

| Trait | Chr | Pos (cM) | LOD | Significance Threshold<sup>†</sup> |
| --- | --- | --- | --- | --- |
| Mn | 1 | 232.4 | 7.03 | 4.59 |
| Mn | 5 | 195.8 | 4.61 | 4.59 |
| Fe | 5 | 204.6 | 4.50 | 3.94 |
| Ni | 1 | 410.3 | 6.15 | 4.69 |
| Ni | 9 | 7.7 | 28.50 | 4.69 |
| Cu | 7 | 165.9 | 5.31 | 4.72 |
| Zn | 4 | 157.4 | 4.44 | 4.13 |
| Rb | 2 | 185.3 | 3.38 | 2.80 |
| Mo | 1 | 378.0 | 48.49 | 4.20 |
| Cd | 2 | 214.6 | 20.26 | 3.87 |

<sup>†</sup> $\alpha = 0.05$

**Figure 2.**
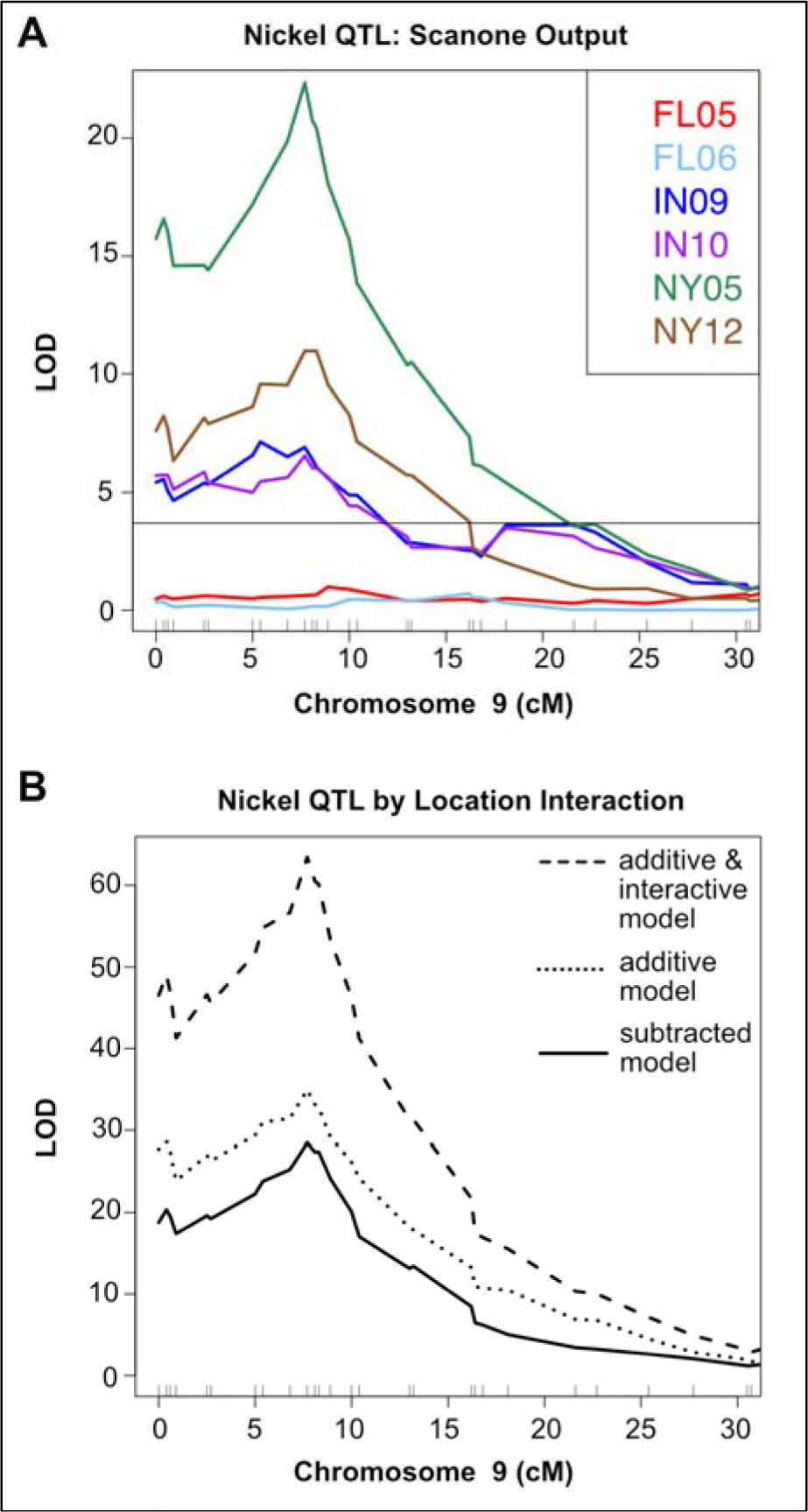
Significant QTL-by-Location Interactions Reflect Variation in Single Environment Mapping. (A) Nickel QTL on chromosome 9 exhibits variation between FL, IN, and NY growouts in single environment QTL mapping. Scanone QTL mapping output for Ni on is plotted for FL05, FL06, IN09, IN10, NY05, and NY12. LOD score is plotted on the y-axis and cM position on the x-axis. Horizontal line corresponds to significance threshold (α = 0.05). (B) Scanone QTL mapping for combined Ni data from Florida (FL05 and FL06), Indiana (IN09 and IN10), and New York (NY05 and NY12) growouts. All lines within each location were included,with covariates designated based on location. QTL mapping output using model with location as an additive covariate is shown as dotted line. QTL mapping output from model with location as both an additive and interactive covariate shown as dashed line. Subtracted LOD score profile from the two models (QTL by location interactive effect only) shown as solid line. Horizontal line corresponds to significance threshold for QTL by location interaction effect, derived from 1000 iterations of the three step procedure using randomized data: scanone QTL mapping with the additive model, scanone QTL mapping with the additive and interactive model, and subtraction of the two models.

## QTL for trait differences within location

The previous method identified genotypes with interactions with location but not with year. Year to year variation will also have effects due to differences in rainfall, temperature and management practices. To examine variation that occurs within a location over different years, we examined the intra-location QEI in the three locations (FL05 & FL06, IN09 & IN10, NY05 & NY12). QTL were mapped using the stepwise algorithm on the trait differences between common lines in the two environments for sample weight and 20 elements. This approach identified loci affecting phenotypic differences between the same lines grown on the same farm but in different years. Six QTL were found for FL05-FL06 differences, one QTL for IN09-IN10 differences, and two QTL for NY05-NY12 differences (Table 3). These trait-difference QTL included locations identified in our single element/environment QTL experiment where a locus was present for one year but not the other or the QTL was found in both years with differing strength (Fig 3A, B, C). Six of the difference QTL were detected at locations where no QTL were detected when the years were mapped separately, revealing novel gene by environment interactions not obvious from the single year data. These significant effects of year to year environmental variation within the same location indicated that factors beyond location are both influencing the ionome and determining the consequences of genetic variation.

**Table 3.** Significant QTL for Trait Differences.

| Location | Years Compared | Trait | Chr | Pos (cM) | LOD | Significance Threshold<sup>†</sup> |
| --- | --- | --- | --- | --- | --- | --- |
| FL | FL05_FL06 | Mg | 8 | 294.4 | 5.23 | 3.74 |
| FL | FL05_FL06 | P | 4 | 130.2 | 3.89 | 3.60 |
| FL | FL05_FL06 | P | 4 | 297.8 | 6.03 | 3.60 |
| FL | FL05_FL06 | P | 8 | 294.6 | 8.43 | 3.60 |
| FL | FL05_FL06 | Co | 1 | 296.3 | 4.36 | 3.69 |
| FL | FL05_FL06 | Mo | 1 | 378.6 | 6.10 | 3.70 |
| IN | IN09_IN10 | Fe | 8 | 140.9 | 4.52 | 3.62 |
| NY | NY05_NY12 | K | 5 | 154.2 | 4.25 | 3.61 |
| NY | NY05_NY12 | Sr | 7 | 193.2 | 4.45 | 3.66 |

**Figure 3.**
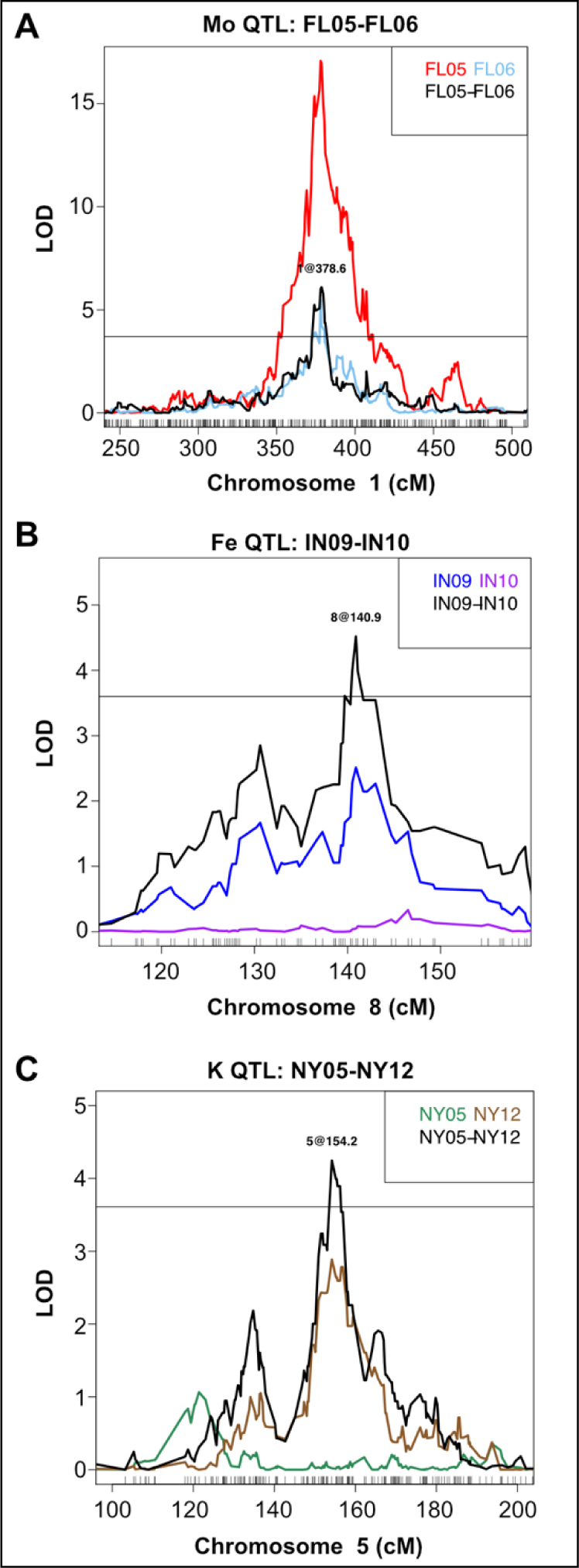
Comparison of QTL Mapped on Traits in Single Environments and TraitDifferences Between Environments. Examples from stepwise QTL mapping on trait differences of between two years at one location, calculated between IBM lines common to both years. Scanone QTL mapping output is plotted for the same trait from each year separately. LOD score is plotted on the y-axis and cM position on the x-axis. Horizontal lines correspond to significance threshold (α = 0.05). (A) Molybdenum QTL on chromosome 1 mapped for Mo in FL05, Mo in FL06, and difference in Mo content between FL05 and FL06. (B) Iron QTL on chromosome 8 mapped for Fe in IN09, Fe in IN10, and difference in Fe content between IN09 h and IN10. (C) Potassium QTL on chromosome 5 mapped for K in NY05, K in NY12, and difference in K content between NY05 and NY12.

### Discussion

The results described here demonstrate that the concentrations of elements in the kernels of maize are strongly affected by the interaction of genetics with growth environment. Dramatically, element concentration is highly heritable within an environment and varied between environments. The presence of a large number of single-environment QTL is consistent with the hypothesis that environment has a significant impact on genetic factors influencing the ionome. By changing the stringency of the statistical tests, we are able to discount the likelihood that that these single environment QTL are the result of a large number of false positives or false negatives. The structure of our data, with few lines measured across all locations and years, limited our ability to test for direct QTL by environment interactions. As a result, we have likely underestimated the extent of QEI. Future studies with uniform lines across environments will allow for inclusion of data from all environments and lines and increase power to detect additional genetic by environment interactions.

Nevertheless, we were able identify QEI over different locations and QEI at a single location over different years. We identified a strong nickel QTL on chromosome 9 that was found in Indiana and New York with single-environment QTL mapping, but not in Florida. This same locus also identified as a significant location-interacting QTL when using a model that included Indiana, New York, and Florida as covariates. One possible cause for this, and other location specific QTL, might be differences in element availability between local soil environments. Interestingly, the presence/absence of the QTL does not seem to correlate with the mean levels of the elements in the grains sampled from that location, suggesting that QEI are not being driven solely by altered availability of the elements in the soil. Local soil differences are less likely to be driving the QTL found for pairwise differences between two years at one location. Soil content should remain relatively similar from year to year at the same farm, suggesting that the loci identified by comparison between years and within location will encode components of elemental regulatory processes responsive to precipitation, temperature, or other weather changes. Experiments with more extensive weather and soil data, or carefully manipulated environmental contrasts, are needed to create models with additional covariates and precisely model environmental impacts.

Although the mapping intervals do not provide gene-level resolution, several QTL overlap with known elemental regulation genes, such as the QTL on chromosome 1 at 378 cM which coincides with ZEAMMB73_045160, an ortholog of the Arabidopsis molybdenum transporter, MOT1. We observe strong effects and replication of this QTL across nearly all environments, suggesting that this MOT1 plays a role in a variety of environments. Other large effect QTL found in several environments merit further investigation, as they may recapitulate important element-associated genes that have yet to be identified. Identification of the genes underlying these QTL and the gene/environmental variable pairs underlying the QEIs will improve our understanding of the factors controlling plant elemental uptake and productivity. Given the high levels of variability that the interaction between genotype and environmental factors can induce in these traits, conventional breeding approaches that look for common responses across many different environments for a single trait may fail to improve the overall elemental content, necessitating rational approaches that include both genetic and environmental factors.

## Conclusions

Here we have shown that the maize kernel ionome is determined by genetic and environmental factors, with a large number of genetic by environment interactions. Elemental profiling of the IBM population across 10 environments allowed us to capture environmentally-driven variation in the ionome. Our QTL analysis on elements found mainly single-environment QTL, indicative of substantial genetic by environment interaction in establishment of the elemental composition of the maize grain. This approach, along with identification of QEI occurring both within a single location over different years and QEI between different locations, demonstrated that gene by environment interactions underlie elemental accumulation in maize kernels.

## Methods

### Field Growth and Data Collection

#### Population and field growth

Subsets of the intermated B73 x Mo17 recombinant inbred (IBM) population were grown in 10 different environments: Homestead, Florida in 2005 (220 lines)and 2006 (118 lines), West Lafayette, Indiana in 2009 (193 lines) and 2010 (168 lines), Clayton, North Carolina in 2006 (197 lines), Poplar Ridge, New York in 2005 (256 lines), 2006 (82 lines),and 2012 (168 lines), Columbia, Missouri in 2006 (97 lines), and Limpopo, South Africa in 2010 (87 lines). In all but three environments, NY05, NC06, and MO06, one replicate was sampled per line. In NY05, 3 replicates of 199 lines, 2 replicates of 50 lines, and 1 replicate of 7 lines were sampled. A replicate is considered pooled ears from a row. Several ears were harvested and kernels were subsampled from pooled ears from the row. After harvesting, seeds were stored in local temperature and humidity controlled seed storage rooms. Subsequently they were shipped to the ionomics lab where they were stored in temperature-controlled conditions. Because each batch of seed was treated identically, any losses in weight or increases in weight due to differing hydration should not affect the relative, weight-adjusted concentrations used for analysis. We do not expect any changes in ion composition due to storage. Table S1 includes planting dates and line numbers after outlier removal and genotype matching. After outlier removal, 199 of the 233 unique lines in the experiment were present in 3 or more of the 10 environments. 106 lines were present in 7 or more of the environments.

### Elemental Profile Analysis

Elemental profile analysis is conducted as a standardized pipeline in the Baxter Lab. The methods used for elemental profile analysis are as described in Ziegler et al. (26). Descriptions taken directly are denoted by quotation marks.

### Sample preparation and digestion

Lines from the IBM population from each environment were analyzed for the concentrations of 20 elements. “Seeds were sorted into 48-well tissue culture plates, one seed per well. A weight for each individual seed was determined using a custom built weighing robot. The weighing robot holds six 48-well plates and maneuvers each well of the plates over a hole which opens onto a 3-place balance. After recording the weight, each seed was deposited using pressurized air into a 16×110 mm borosilicate glass test tube for digestion. The weighing robot can automatically weigh 288 seeds in approximately 1.5 hours with little user intervention.”

“Seeds were digested in 2.5 mL concentrated nitric acid (AR Select Grade, VWR) with internal standard added (20 ppb In, BDH Aristar Plus). Seeds were soaked at room temperature overnight, then heated to 105°C for two hours. After cooling, the samples were diluted to 10 mL using ultrapure 18.2 MΩ water (UPW) from a Milli-Q system (Millipore). Samples were stirred with a custom-built stirring rod assembly, which uses plastic stirring rods to stir 60 test tubes at a _time. Between uses, the stirring rod assembly was soaked in a 10% HNO_3 solution. A second dilution of 0.9 mL of the 1st dilution and 4.1 mL UPW was prepared in a second set of test tubes. After stirring, 1.2 mL of the second dilution was loaded into 96 well autosampler trays.”

### Ion Coupled plasma mass spectrometry analysis

Elemental concentrations of B, Na, Mg, Al, P, S, K, Ca, Mn, Fe, Co, Ni, Cu, Zn, As, Se, Rb, Sr, Mo, and Cd “were measured using an Elan 6000 DRC-e mass spectrometer (Perkin-Elmer SCIEX) connected to a PFA microflow nebulizer (Elemental Scientific) and Apex HF desolvator (Elemental Scientific). Samples were introduced using a SC-FAST sample introduction system and SC4-DX autosampler (Elemental Scientific) that holds six 96-well trays (576 samples).” Measurements were taken with dynamic reaction cell (DRC) collision mode off. “Before each run, the lens voltage and nebulizer gas flow rate of the ICP-MS were optimized for maximum Indium signal intensity (>25,000 counts per second) while also maintaining low CeO+/Ce+ (<0.008) and Ba++/Ba+ (<0.1) ratios. This ensures a strong signal while also reducing the interferences caused by polyatomic and double-charged species. Before each run a calibration curve was obtained by analyzing six dilutions of a multi-element stock solution made from a mixture of single-element stock standards (Ultra Scientific). In addition, to correct for machine drift both during a single run and between runs, a control solution was run every tenth sample. The control solution is a bulk mixture of the remaining sample from the second dilution. Using bulked samples ensured that our controls were perfectly matrix matched and contained the same elemental concentrations as our samples, so that any drift due to the sample matrix would be reflected in drift in our controls. The same control mixture was used for every ICP-MS run in the project so that run-to-run variation could be corrected. A run of 576 samples took approximately 33 hours with no user intervention. The time required for cleaning of the instrument and sample tubes as well as the digestions and transfers necessary to set up the run limit the throughput to three 576 sample runs per week.”

## Computational Analysis

### Drift correction and analytical outlier removal

Analytical outliers were removed from single-seed measurements using a method described in Davies and Gather (1993). Briefly, values were considered an outlier and removed from further analysis if the median absolute deviation (MAD), calculated based on the line and location where the seed was grown, was greater than 6.2.

Normalization for seed weight by simply dividing each seed’s solution concentration by sample weight resulted in a bias where smaller seeds often exhibited a higher apparent elemental concentration, especially for elements whose concentration is at or near the method detection limit. This bias is likely either a result of contamination during sample processing, a systematic over or under reporting of elemental concentrations by the ICP-MS or a violation of the underlying assumption that elemental concentration in seeds scales linearly with seed weight. Instead, we developed a method whereby the residuals from the following linear model:

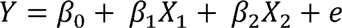

where Y is the non-weight normalized measure of elemental concentration for each seed after digestion, β_0_ is the population mean, *X*_1_ is the seed weight, *X*_2_ is the analytical experiment the seed was run in (to further correct for run-to-run variation between analytical experiments), and *e* is the residual (error) term. The residuals in this linear model represent how far each data point departs from our assumption that analyte concentration will scale linearly with sample weight. If all samples have the same analyte concentration then the linear model will be able to perfectly predict analyte concentration from weight and the residuals will all equal zero. However, if a sample has a higher or lower concentration of an analyte then the general population being measured, then it will have a residual whose value represents the estimated concentration difference from the population mean. For this reason, we have termed this value the estimated concentration difference from the mean (ECDM).

### Heritability calculation

Broad-sense heritability was calculated for seed weight and elements across environments and within three environments for which we had substantial replicate data. To calculate the broad-sense heritability across 10 environments, the total phenotypic variance was partitioned into genetic and environmental variance, with the broad-sense heritability being the fraction of phenotypic variance that is genetic. This was done using an unbalanced, type II analysis of variance (ANOVA) in order to account for the unbalanced common line combinations across environments. Two models were fit using the *lmfit* function in R. The first model included genetic variance as the first term and environmental variance as the second. The second model had the opposite form. The sum of squares for genetic or environmental components was obtained using the *anova* function on the model in which that component was the second term. Broad-sense heritability was calculated by dividing the genetic sum of squares by the total (genetic plus environmental) sum of squares. Heritability was calculated within environments for NY05, NC06, and MO06. Data with outliers designated as NA was used for each environment. For each element within an environment, lines with NA were removed and lines with only 1 replicate were removed, leaving only lines with 2 or more replicates. The heritability was then calculated for seed weight and each element using *lmfit* followed by *anova* functions to obtain the sum of squares for the genetic component and the residuals. Broad-sense heritability was calculated as the proportion of total variance (genetic plus residuals) explained by the genetic component.

### QTL mapping: elemental traits

The R package R/qtl was used for QTL mapping. For each of the 10 environments, elemental trait line averages and genotypes for all lines, 4,217 biallelic single nucleotide polymorphisms (SNPs) distributed across all 10 maize chromosomes, were formatted into an R/qtl cross object. The *stepwiseqtl* function was used to implement the stepwise method of QTL model selection for 21 phenotypes (seed weight, 20 elements). The max number of QTL allowed for each trait was set at 10 and the penalty for addition of QTL was set as the 95th percentile LOD score from 1000 *scanone* permutations, with imputation as the selected model for *scanone*. A solely additive model was used; epistatic and interaction effects were not considered and thus heavy and light interaction penalties were set at 0. QTL positions were optimized using *refineqtl*, which considers each QTL one at a time, in random order, iteratively scanning in order to move the QTL to the highest likelihood position. QTL models for each trait in each environment were obtained using this procedure. QTL within 5 cM of each other were designated as the same QTL.

### QTL by environment analysis: linear model comparison

Linear modeling was used determine instances and strength of QEI using all data from two years within three locations (FL, IN, NY). The specific growouts analyzed together were FL05, FL06, IN09, IN10, NY05, and NY12. FL, IN, and NY were then used as covariates in QTL analysis. Two QTL models, one with location as an additive and interactive covariate and one with location as only an additive covariate, were fit for each phenotype (sample weight, 20 elements) using the *scanone* function in R/qtl,

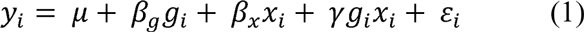

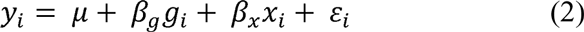

where *y_i_* is the phenotype of individual *i*, *g_i_* is the genotype of individual *i*, and *x_i_* is the location of individual *i*. *B_g_* and *B_x_* are additive effects of genotype and environment, respectively, and γ is the effect of genotype by environmental interaction. LOD scores for each marker using model (2) were subtracted from LOD scores for each marker using model (1) to the isolate genetic by location effect. QTL by location interaction was determined as QTL with a significant LOD score after subtraction. The significance threshold was calculated from 1000 permutations of the three step procedure (fitting the two models and then subtracting LOD scores) and taking the 95th percentile of the highest LOD score.

### QTL by environment analysis: mapping on within-location differences

QTL were mapped on phenotypic differences between common lines grown over two years at a single location. This procedure was used to compare FL05 and FL06, IN09 and IN10, and NY05 and NY12 by calculating the differences for each trait value between common lines in location pairs (FL05-FL06, IN09-IN10, NY05-NY12) and using these differences for analysis using the previously described *stepwiseqtl* mapping and permutation procedure.

## Acknowledgements

The authors would like to thank Justin Borevitz and Riyan Cheng, and Tom Juenger for advice on QTL by environment mapping and Karl Broman for invaluable assistance with R/qtl. The authors would especially like to thank our field collaborators Sherry Flint-Garcia, Peter Balint-Kurti, Torbert Rocheford, Jonathan Lynch, and Robert Snyder for their dedicated efforts to provide the seeds analyzed for this project.

## Supporting Information

S1 Table. Growout Information.

S2 Table. Percent Variance (R2) of Mo, Cd, and Ni QTL.

S3 Table. Location LOD Scores Compared to Seed Element Content.

